# Lightweight bioinformatics: evaluating the utility of Single Board Computer (SBC) clusters for portable, scalable Real-Time Bioinformatics in fieldwork environments via benchmarking

**DOI:** 10.1101/337212

**Authors:** Joe Parker

## Abstract

The versatility of the current DNA sequencing platforms and the development of portable, nanopore sequencers means that it has never been easier to collect genetic data for unknown sample ID. In fact, the distinction between fieldwork and the laboratory is becoming blurred since genome-scale data can now be collected in challenging conditions in a matter of hours. However, the full scientific and societal benefits of these new methods can only be realised with equally rapid and portable analyses. At present, field-based analyses of genomic data, despite advances in computing technology, remain problematic; laptop computers are relatively expensive and limited in scalability, while cloud- and cluster-based analyses depend, for the time being, on sufficiently reliable high-bandwidth data uplinks to transmit primary data for analysis.

Single board computers (SBCs), such as the Raspberry Pi, offer a potential solution to this problem: while less powerful than their laptop cousins, their very individual low cost and power consumption mean modest arrays of SBCs could be used for field-based preprocessing, or complete analyses or primary data. In this study we investigate the performance of one SBC, the Pi 3 Model B+, on a range of typical field-sequencing tasks versus laptop and cloud-based form-factors. Our data analysis pipeline has been made available as a workflow on Github for simple, scalable deployment for a range of uses.

## Introduction

The nucleic acids present in every living organism provide not only *the* essential information needed for species identification to characterise biodiversity in a sample, site, or ecosystem. They can also be interpreted to reconstruct evolutionary histories hundreds of millions of years back into geologic time, or furnish us with detailed information about the metabolic and immunological activity of a sample of mud, blood, or water. From the discovery of the structure of DNA (1953) this information was hard-won, requiring laboratories with specialised equipment and staff.

This is changing, fast. Portable single molecule, real-time DNA sequencers (such as the Oxford Nanopore MinION have now become a commercial reality. Portable sequencers allow DNA-sequencing to happen anywhere, in real-time, with important applications that include disease surveillance and food-chain monitoring. In less than a decade, these devices have moved from the drawing-board to the mainstream of genomics and there is every suggestion that science is set to undergo a transformation: millions of researchers, clinicians, conservation professionals and citizen-scientists will have the potential to sequence and analyse genomic material anytime, anywhere (Erlich, 2015). Uses so far have included epidemic monitoring in Guinea (Quick *et al.,* 2016), extremophile sequencing in Antarctica (Michael *et al.*, 2017), assembly of complete plant genomes on a single flowcell, in a week (Johnson *et al.*, 2017), species ID and phylogenomics in the field (Parker *et al.*, 2017, 2018). The real-time nature in which reads are generated, and the very long length of nanopore reads compared with traditional high-throughput sequencing (HTS) inserts (tens of thousands of base-pairs compared with a few hundred), plus the availability of PCR-free direct sequencing methods, also make multilocus metagenomics and phylogenomics possible; a potential advantage over molecular barcoding approaches, which are far slower and also subject to error arising from reticulate (non-tree-like) evolution which can confound identification and inference (Mallo & Posada 2016; Liu et al. 2017).

While bioinformaticians and experimentalists are well-versed in the seeming existence of Moore’s Law as applicable to high-performance computing (HPC) clusters and its implications, the past decade has also seen a parallel rise in the availability and interest of single-board computers (SBCs), most notably the Raspberry Pi family. These stripped-back computing devices are cheap (€50 or so), tiny – typically described as ‘credit-card-sized’, although many are smaller still – and typically consume a tenth of the power needed for a laptop, much less a powerful desktop, fatnode or cluster. Several authors have noted the potential for small (4-20 node) SBC clusters to replace on-site laptops or remote computing resources, in diverse applications ranging from cyberwarfare (Matthews, 2016) to teaching (Barker *et al*, 2013; Cox *et al.*, 2013). The application of SBC clusters to field-sequencing is nonetheless as immature as the field-sequencing devices are new.

In the present study, we investigate the utility of SBCs for field-based bioinformatics analyses. Several analyses of previously published datasets are carried out to test the performance of the Raspberry Pi 3, perhaps the most common SBC available at the present time, in carrying out typical tasks. Run times are benchmarked and compared to typical values for other platforms. Finally, the wider context and possible future development of this field are considered.

## Methods

### Equipment

To conduct the present study we assembled a small cluster (6 nodes, plus headnode; see Figure 1a) of Raspberry Pi 3 Model B+ SBCs, and used this cluster to benchmark typical field-sequencing genomics analyses tasks in comparison to a consumer laptop (Apple MacBook Pro, 2011) and a bioinformatics fatnode / enterprise HPC machine (Dell PowerEdge). Field-analysis cluster components are listed in Table 1; specifications for comparison machines in Table 2.

**Table 1:**
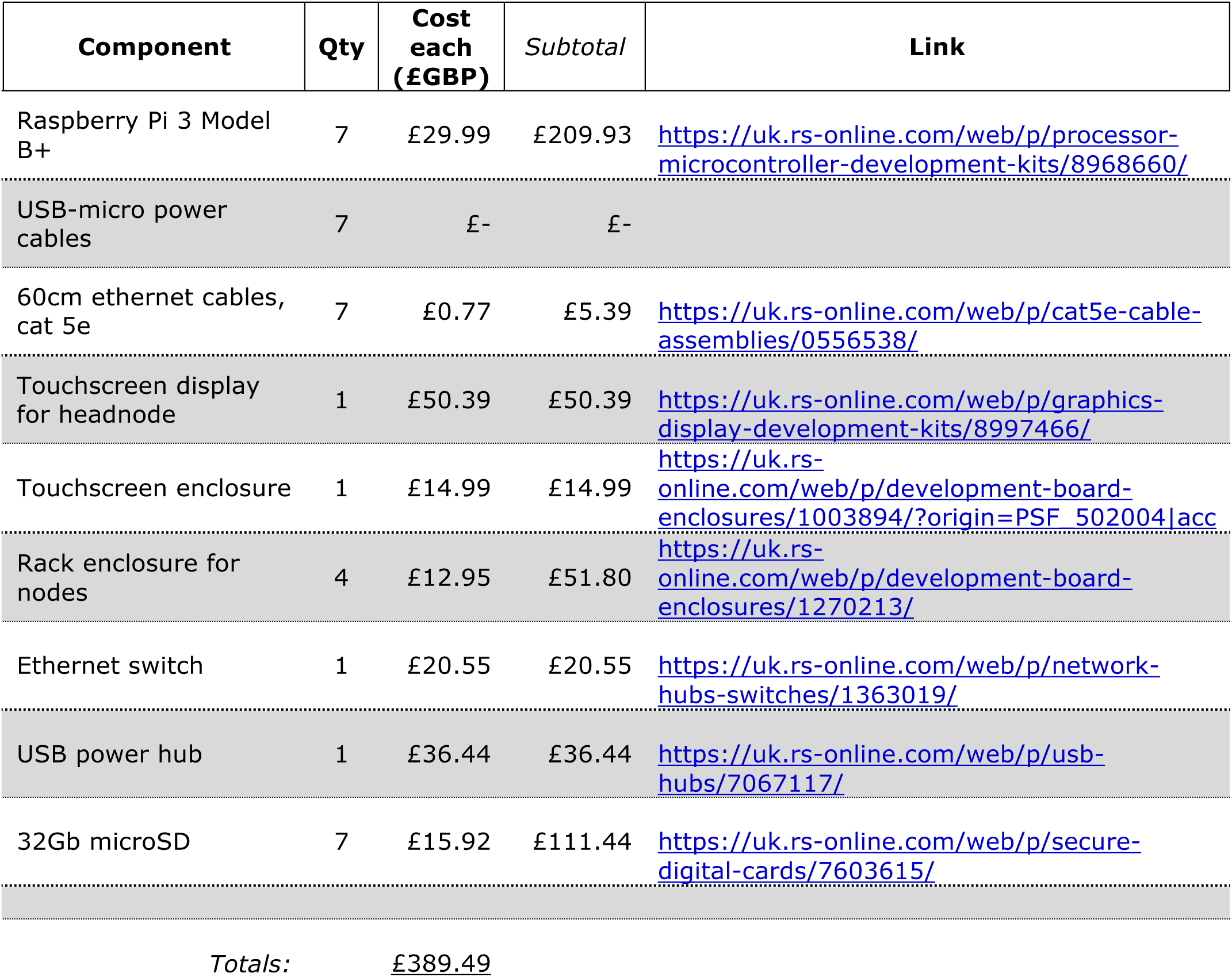
Component list for Raspi field-sequencing cluster

**Table 2:** Comparison of systems evaluated in this study.

| System | Architecture | CPU type, clock GHz | Number of cores | RAM Gb | Scratch size, Gb |
| --- | --- | --- | --- | --- | --- |
| Lab node (PowerEdge) | i686 | Xeon E5620, 2.4 | 8 | 64 | 1000, SSD |
| Raspberry Pi 2 B+ | ARM | ARMv7, 1.0 | 1 | 1 | 8, microSD |
| Raspberry Pi 3 B+ | ARM | ARMv7, 1.2 | 1 | 1 | 32 microSD |
| Macbook Pro (2011) | x64 | Core i7, 2.2 | 4 | 8 | 250, SSD |
| (EC2 m4.10xlarge ) | x64 | Xeon E5, 2.4 | 40 | 160 | 320, SSD |

**Figure 1a:**
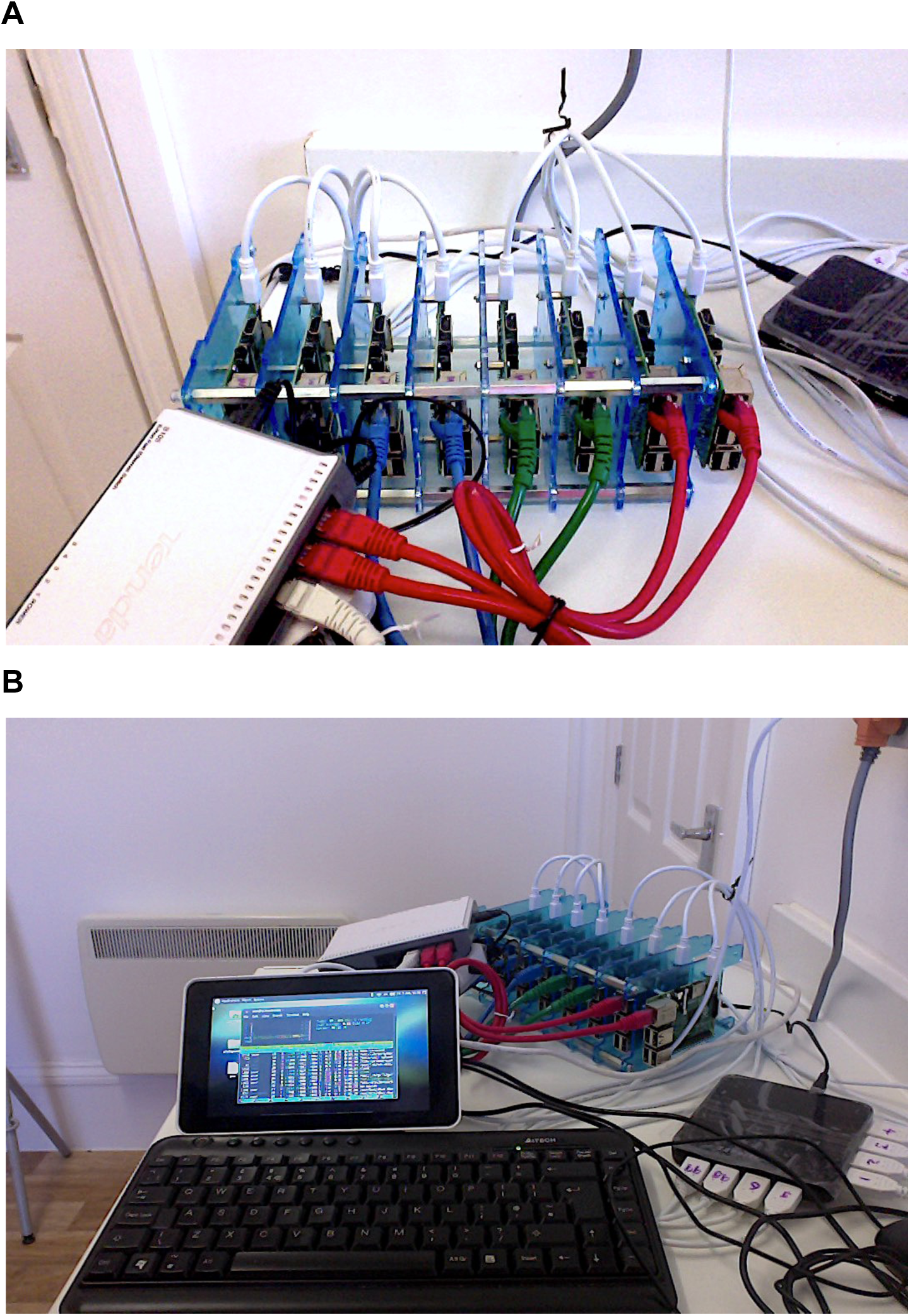
Picture of bioinformatics cluster for field-sequencing comprising 8 Raspberry Pi SBCs plus headnode. *A*, detail of worker nodes, enclosure and cableing; *B*, overview of the whole cluster, including headnode with IO peripherals. The cluster (excluding power) fits into a standard daysack.

### Data

Source data was taken from field-sequencing studies previously reported in Parker *et al.* (2017) and Parker *et al.* (2018a); see Table 3. Briefly DNA was extracted from fresh plant tissue using commercial kits (Qiagen DNEasy Plant Miniprep) and whole genome shotgun libraries were prepared for MinION R9 and R9.5 chemistry using rapid (SQK-RAD001/RAD003) protocols and kits. Field-extraction and sequencing were carried out in Richmond Park, London, and Snowdonia National Park, Wales. A full list of equipment required for field-sequencing is given in the Supplementary Information for those papers.

**Table 3:** Summary BLASTN results. Reads from the *A. thaliana* dataset were subsampled without replacement and matched to the TAIR10 genome database with BLASTN.

| Machine | Number of threads | Number of queries | Mean wall clock time (s) | Std. dev |
| --- | --- | --- | --- | --- |
| Lab node | 1 | 10 | **0.17** | 0.11 |
|  |  | 100 | **1.12** | 0.84 |
|  |  | 1000 | **8.18** | 3.18 |
|  |  | 10000 | **58.67** | 7.53 |
|  | 2 | 10 | **0.14** | 0.03 |
|  |  | 100 | **0.75** | 0.38 |
|  |  | 1000 | **3.98** | 2.04 |
|  |  | 10000 | **41.70** | 3.53 |
|  | 4 | 10 | **0.11** | 0.02 |
|  |  | 100 | **0.71** | 0.41 |
|  |  | 1000 | **2.96** | 0.39 |
|  |  | 10000 | **31.38** | 1.40 |
|  | 8 | 10 | **0.09** | 0.00 |
|  |  | 100 | **1.43** | 1.91 |
|  |  | 1000 | **2.93** | 1.67 |
|  |  | 10000 | **34.10** | 2.64 |
| MacBook Pro | 1 | 10 | **0.53** | 0.07 |
|  |  | 100 | **2.23** | 0.93 |
|  |  | 1000 | **20.43** | 4.66 |
|  | 2 | 10 | **0.48** | 0.09 |
|  |  | 100 | **1.56** | 0.57 |
|  |  | 1000 | **23.45** | 13.69 |
|  | 4 | 10 | **0.44** | 0.12 |
|  |  | 100 | **1.87** | 1.43 |
|  |  | 1000 | **17.09** | 8.48 |
|  | 8 | 10 | **0.59** | 0.21 |
|  |  | 100 | **2.87** | 2.11 |
|  |  | 1000 | **16.73** | 9.89 |
| Raspberry Pi 2 | 4 | 10 | **3.04** | 1.09 |
|  |  | 50 | **6.77** | 5.08 |
|  |  | 100 | **11.79** | 9.23 |
|  |  | 500 | **33.68** | 17.65 |
|  |  | 1000 | **56.52** | 20.25 |
|  |  | 5000 | **262.25** | 40.06 |
| Raspberry Pi 3 | 1 | 10 | **37.62** | 49.64 |
|  |  | 100 | **18.86** | 1.96 |
|  | 2 | 10 | **9.06** | 1.67 |
|  |  | 100 | **14.47** | 2.13 |
|  | 3 | 10 | **11.37** | 2.96 |
|  |  | 100 | **19.32** | 9.87 |
|  | 4 | 10 | **3.16** | 2.16 |
|  |  | 50 | **9.08** | 15.92 |
|  |  | 100 | **10.12** | 6.92 |
|  |  | 500 | **31.39** | 17.73 |
|  |  | 1000 | **49.17** | 19.47 |
|  |  | 5000 | **241.11** | 40.32 |

### Operating system/pipeline

Ubuntu 16.04.3 LTS was installed on all nodes. Having experimented with various job schedulers (Condor, Slurm) and chunking approaches, we determined that the most stable configuration was also the simplest (nodes dedicated to individual tasks, working in series from a common read pool). A watch-script was used to monitor a shared 1TB NFS drive, to which the MinION sequencing laptop also had write access to deposit newly sequenced reads in real-time. In prototype, 1000-read chunks were analysed and moved sequentially through the pipeline (shown in Figure 1b).

### Guppy basecaller

The basecaller is an algorithm responsible for converting raw measurements from the sequencing machine into nucleotide sequences. We used the experimental Guppy basecaller, source provided by Oxford Nanopore Technologies. To benchmark performance a complete set of 4000 reads (N50 1.9kbp; max >50kbp) from the Parker *et al.* (2018b) acute oak decline study was analysed for five replicates, and an approximate handling time per read averaged.

### Read mapping/matching

BLASTN (Camacho 2008) is typically used to test all reads for the presence of host, contaminant/control (human; phage lambda) and target pathogen DNA sequences. In benchmarking, reads from the 2017 Parker *et al.* study were subsampled randomly without replacement (as in Parker *et al.*, 2018a) and matched against the *A. thaliana re*ference genome (TAIR10) using BLAST with 1, 2, 3 or 4 threads and only best hits retained.

### ab initio gene prediction

SNAP was used to predict the occurrence of coding genes directly from individual reads.

### Phylogeny inference

Once a set of orthologous genes have been identified, they can be aligned and a phylogeny (evolutionary hypothesis) inferred. Multiple alignments comprising 6-10 taxa and 500-2000bp were created from theten best SNAP-identified, BLAST-checked genes with Muscle and phylogenies inferred with RAxML (as in Parker *et al.,* 2017)

### Metagenomic classification

Kraken (Wood, 2014) were used for metagenomic classification of MinION reads from mixed samples using *k*-mer hashing, here tested against the acute oak decline dataset.

## Results

### Guppy basecaller

4000 .fast5 reads were basecalled using Guppy. Results are given in Table S1. To execute adequately, the options for single threading and small (1000) chunk size were required. Basecalling the whole set on a Raspberry pi required mean execution real / user time of 15,146/30,217s (s.d., 213/426s) for the set (approx 3.5s (real) / 7.6s (user) per read; *N*=5). On the fatnode mean execution real / user time was 530/3194s (s.d., 33/27s) for the set (approx 0.25s (real) / 1s (user) per read; *N*=3). Effectively, reads were basecalled seven times slower using a single Pi node.

### Read mapping/matching

BLASTN (Camacho 2008) ran adequately on the Pi SBCs by comparison to other systems running equivalent query task sizes (Table 3; Table S2; Figure 3a). As expected, wall clock time increased approximately as *O*(n) with number of reads, with each platform analysed (Figure 3b).

**Figure 2b:**
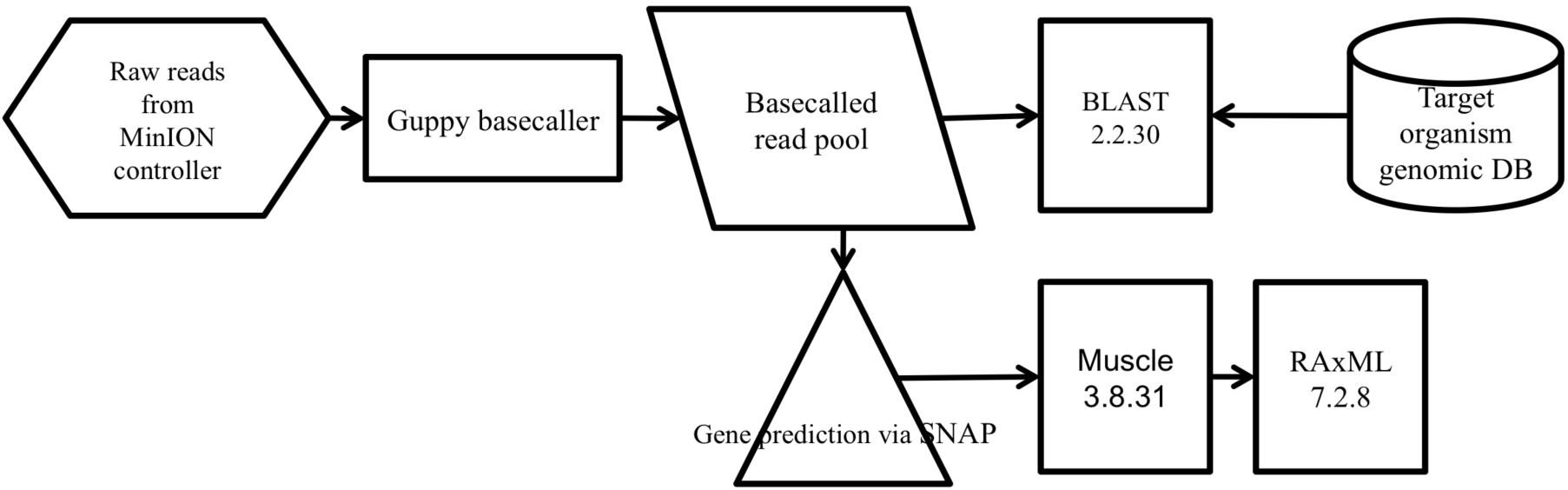
Block diagram of principal workflow units. Reads are passed from the MinION controller (laptop running MinKNOW client) to the common read pool (shared NFS drive on SBC cluster). Individual nodes are assigned discrete tasks in the pipeline (delimited by subdirectory structure) with responsibility for monitoring upstream progress and passing completed outputs to the next node.,

**Figure 3:**
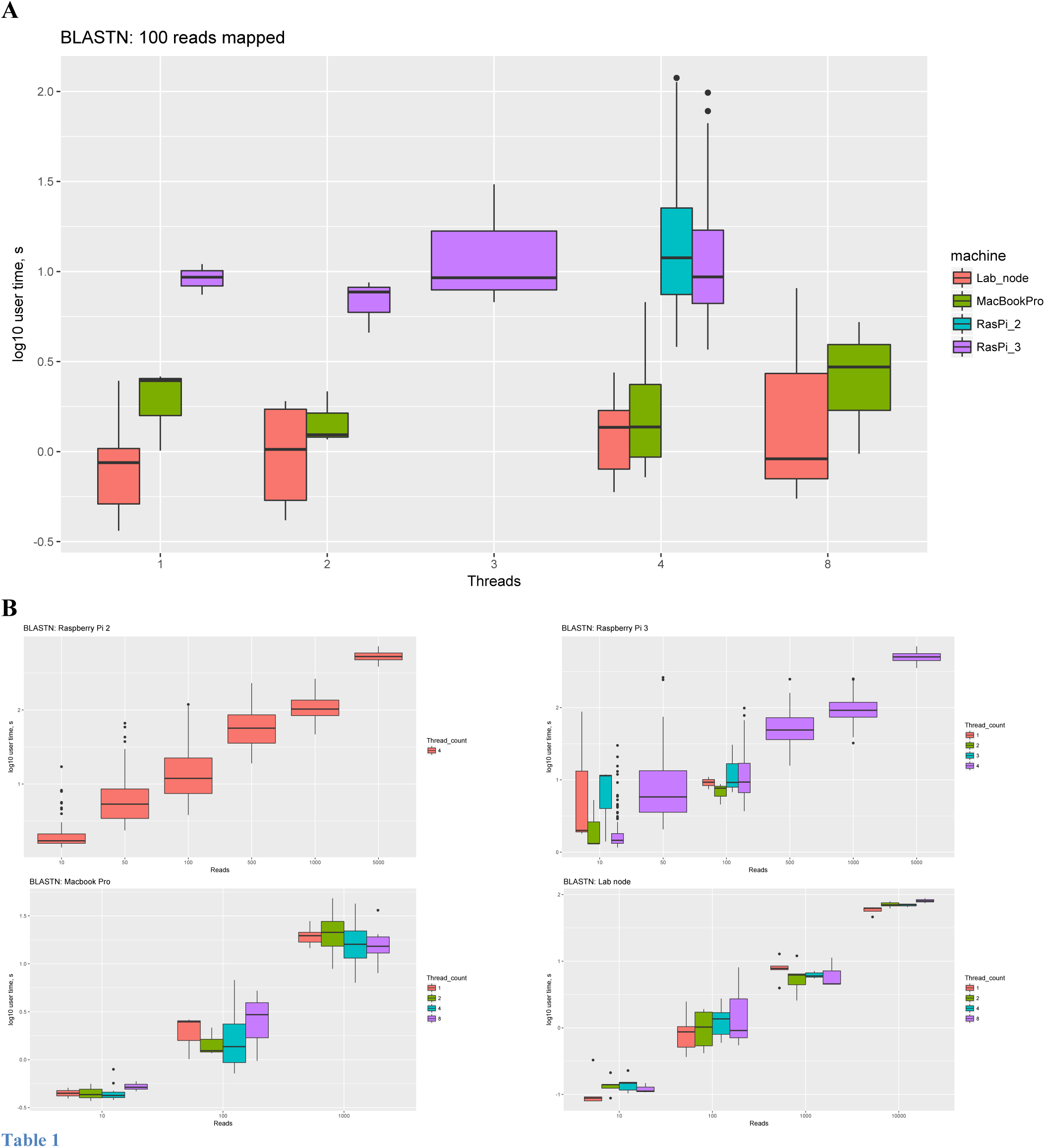
Performance of single cluster node on sample classification using BLASTN. A, log_10_ user time (seconds) to map 100 reads, mean of 30 replicates. B, performance of (clockwise from top-left): Raspberry Pi 2; Raspberry Pi 3; Lab node; Macbook Pro.

### ab initio gene prediction, alignment, and phylogeny inference

SNAP ran adequately on all systems and installed simply on the Pi’s ARM architecture. However execution as measured by wall-clock time was approximately an order of magnitude slower than for the lab node (Figure 4a; Table S3). Similarly, **m**uscle and RAxMLboth installed to the Pi easily, and in series ran well on the SBC nodes though an order of magnitude slower than the lab node (Table S4; Figure 4b). Surprisingly higher thread count did not make an appreciable difference to run times.

**Figure 4a:**
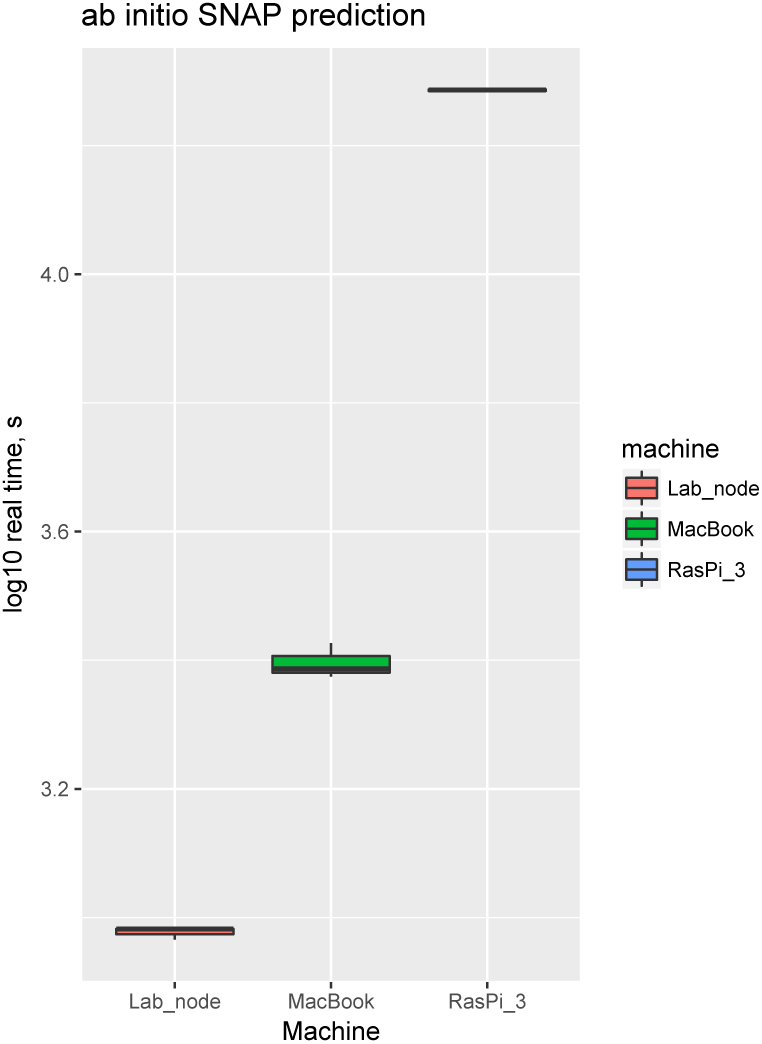
Performance of single cluster node on direct raw-read annotation using SNAP. 96,000 Nanopore reads were annotated using SNAP (*N*=5).

**Figure 4b:**
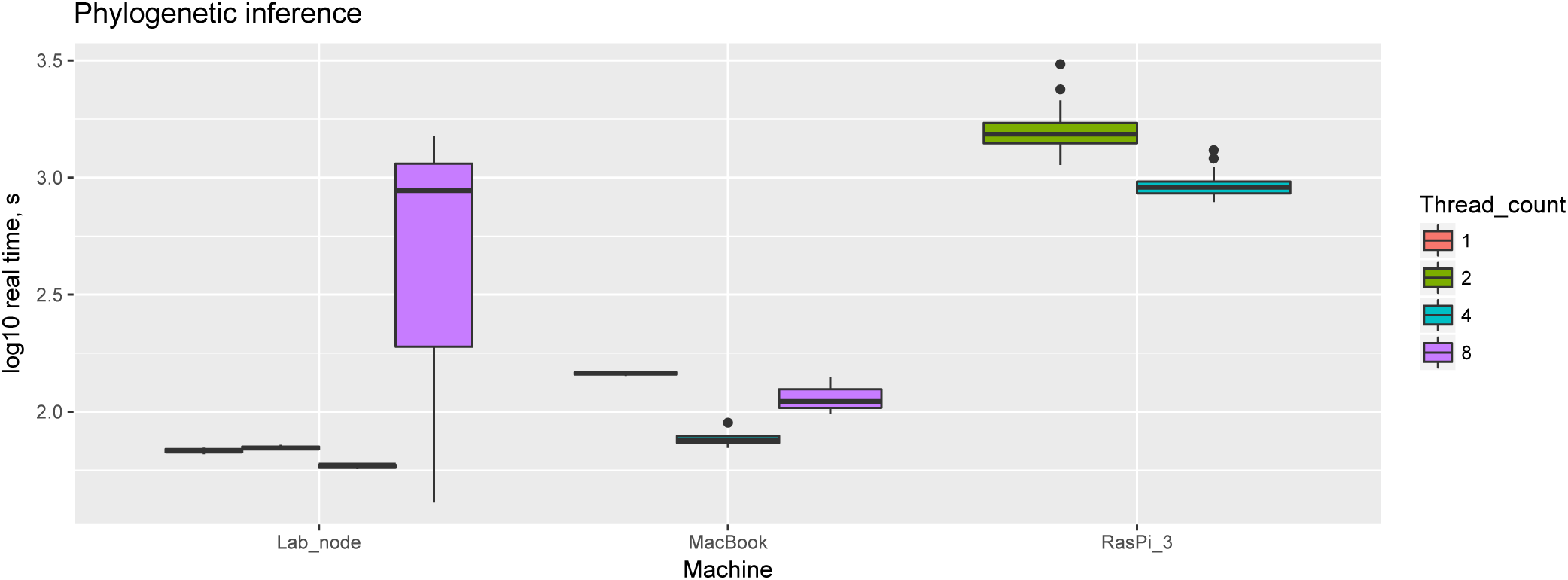
Performance of single cluster node on phylogeny inference using Muscle and RAxML. A dataset of 10 coding genes was aligned and a phylogeny inferred with varying numbers of threads (*N*=5)

### Metagenomic classification

Kraken metagenomic classification of reads from mixed samples did not perform well. Owing to RAM constraints, only the very smallest databases could be queried.

## Discussion

### Successes: high-compute, low-memory tasks (BLASTN, Muscle, RAxML)

Those tasks focused on CPU resource rather than memory performed well on the SBCs, even in comparison with the other systems (roughly as a function of clock speed, unsurprisingly).

### Plausible: high-compute, intermediate-memory tasks (Guppy basecaller)

We found that execution of the experimental Guppy basecaller was plausible for these machines, but higher memory (RAM) constraints meant the low availability on these SBCs limited performance, and careful argument optimisation was needed to gain stable behaviour. Nonetheless, since performance on a single node was within an order of magnitude to that obtained with a lab node, it is feasible that a larger number of Pis (perhaps 4-8) could keep pace with real-time nanopore read generation easily.

### A failure: metagenomic classification via Kraken (high-memory)

Kraken is usually recommended for a minumum of 8Gb RAM and unsurprisingly all but the smallest databases (a few taxa) could not be loaded into the limited physical RAM (1Gb) found on the Raspberry Pis.

### Advantages of SBC clusters

Aside from their low cost (and so scalability), the power consumption and portability of SBC clusters compares well with other systems; a grid of 10-20 SBCs, powered from a single AC generator outlet or vehicle cell, would draw no more than 10A/50W (∼0.5A, 5vDC each), comparable to (or less than) a single laptop, and with more computational power. A 20-node SBC cluster could, with careful design, dissemble into a small rucksack.

### Outstanding challenges

The main shortcoming of these systems is their low RAM, since this precludes the Kraken metagenomic classifier (and genome assembly). However, we have previously argued (Parker *et al.* 2017; 2018a) that classification of extremely long, but noisy, reads using an exact *k*-mer approach (as in Kraken) is counterintuitive, and shown that mapping whole reads (BLASTN; Exonerate; LASTAL) has many underappreciated merits; in this context, the good performance of BLASTN on the SBC cluster is heartening. Our job scheduling/load balancing approach is also naï ve and while full MPI parallelisation is unlikely to be efficient (given the limited node interconnect bandwidth available for these systems), further work to optimise an existing scheduler deployment (Slurm; Condor) or devise a new reatltime system, perhaps based on Watchdog or node.js, is likely to yield quick rewards. Finally it should be borne in mind that, since these clusters’ main use is envisaged for low-bandwidth sites, pushing software updates or expanding reference datasets in the field will remain challenging.

## Conclusion

We have shown that SBC clusters are adequate for a surprising range of useful bioinformatics tasks related to field-based DNA sequencing and analysis. While a single SBC’s performance is inferior to a laptop (let alone an HPC/cloud resource) in every aspect except power consumption, the performance gap is not too great to render SBC clusters adequate to perform analyses in cases where cost is a factor, an expensive laptop might not survive, or insufficient bandwidth exists for uplink to remote resources. However, high-memory tasks, including *de novo* assembly, remain outside the scope of these architectures, and are likely to remain so for the near-future. In addition, improved load balancing / job scheduling efficiency for these resources would greatly improve their utility.

## Acknowledgements

The author thanks James Crowe at RBG Kew, Dan Barker at the University of St. Andrews, Simon Cox at the University of Southampton and Suzanne J. Matthews at USMA for advice and encouragement. This work was funded by a Pilot Study Fund award to JDP from the Kew Foundation.

